# Integrative multi-omics and multi-trait analysis prioritizes regulatory mechanisms and genes for metabolic dysfunction-associated steatotic liver disease

**DOI:** 10.64898/2026.02.19.706502

**Authors:** Zhaode Feng, Fangjun Chen, Jiajia Xiao, Anhui Du, Jianmin Deng, Sirui Wu, Yi Zhang, Xiaokang Li, Adi Zheng, Hao Li

## Abstract

Metabolic dysfunction-associated steatotic liver disease (MASLD) is a prevalent condition that progresses from simple steatosis to advanced fibrosis, significantly affecting liver function and systemic health. Despite its widespread impact, therapeutic options are limited, highlighting the urgent need for comprehensive exploration to identify potential therapeutic targets. In this study, we created an analysis pipeline anchored on liver gene expression, integrating differential meta-analysis of transcriptomic data across three MASLD stages, transcriptome-wide Mendelian randomization (MR), and transcriptome-wide association studies (TWAS), to identify 39 candidate genes potentially involved in MASLD progression. Furthermore, we prioritized these genes using a scoring system that incorporated gene expression-clinical phenotype correlation meta-analysis, proteome-wide association studies (PWAS), and external genetic data from the GWAS Catalog and ExPheWAS. Single-nucleus RNA sequencing (snRNA-seq) analysis of liver cells from healthy to cirrhotic stages revealed stage- and cell-type-specific expression patterns of these prioritized genes. Through experimental validation in a lipid overload hepatocyte model, we confirmed the role of *MLIP* in lipid metabolism. These findings, available through an interactive web portal (masldportal.net), provide valuable insights into MASLD mechanisms and offer an easy-accessible resource for the research community.

**Graphic abstract:** 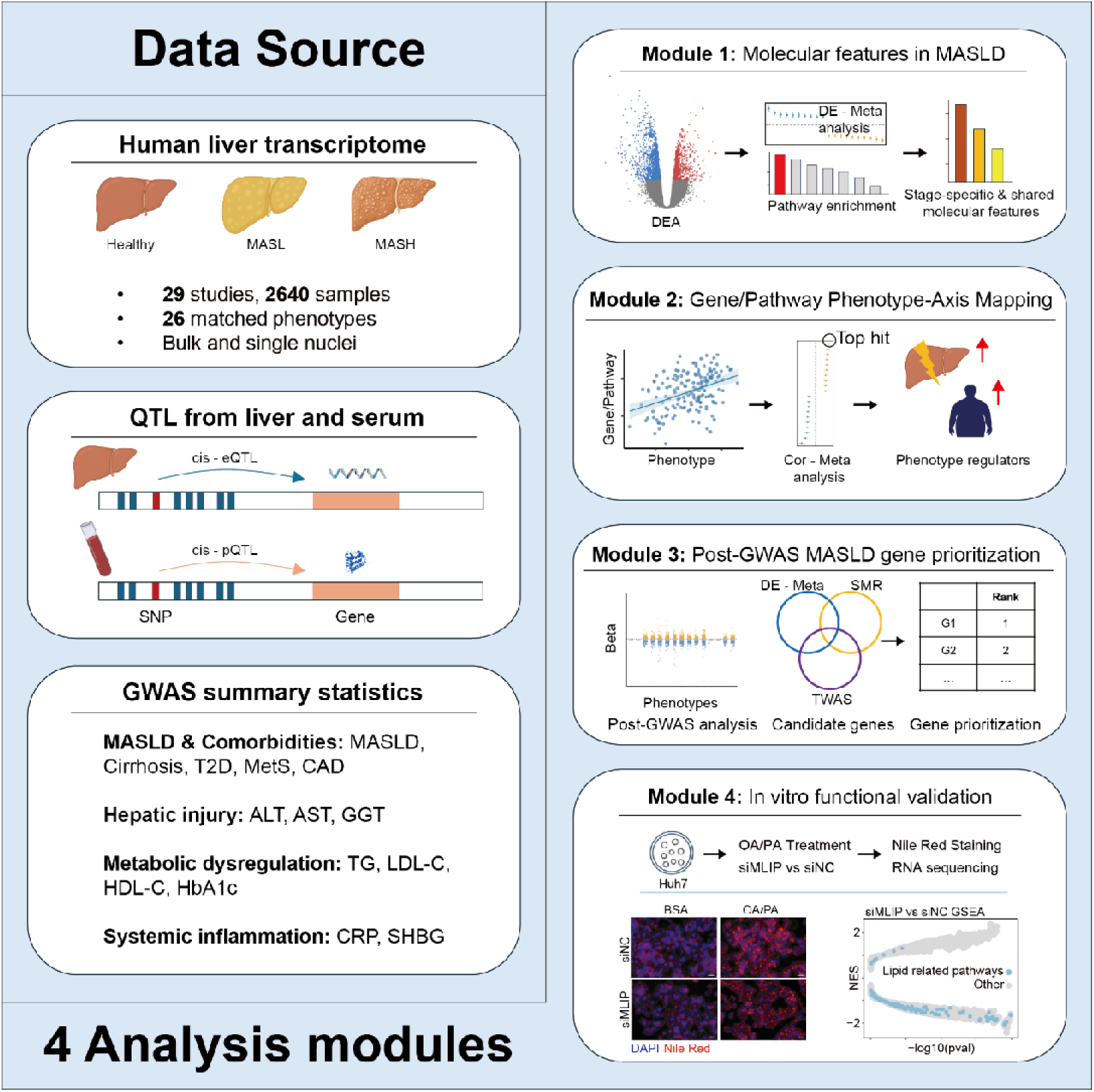

## Introduction

Metabolic dysfunction–associated steatotic liver disease (MASLD), formerly termed non-alcoholic fatty liver disease (NAFLD), is now the most prevalent chronic liver disorder worldwide, affecting an estimated 38% of adults and 7–14% of children, with prevalence projected to exceed 55% by 2040^1^. MASLD encompasses a histological spectrum from steatosis (MASL) to metabolic dysfunction–associated steatohepatitis (MASH) and progressive fibrosis (F0–F4). Across this continuum, fibrosis stage is the strongest determinant of prognosis, with advancing fibrosis linked to a steep increase in all-cause mortality and liver-related death^2^. Clinically, MASLD has become a leading indication for liver transplantation and an increasingly important driver of hepatocellular carcinoma—including in some non-cirrhotic livers—and it substantially elevates the risk of cardiovascular disease, chronic kidney disease, type 2 diabetes, and several extra-hepatic cancers, underscoring its multisystem nature^3^.

Therapeutic options remain limited. Till now, only two agents have received accelerated FDA approval for noncirrhotic MASH with F2-F3 fibrosis: the THR-β agonist resmetirom (MASH resolution without worsening of fibrosis, ∼26–30%; fibrosis improvement without worsening of MASH, ∼24–26%)^4,5^ and semaglutide 2.4 mg (Wegovy) (MASH resolution without worsening of fibrosis, 62.9%; fibrosis improvement without worsening of MASH, 36.8% at week 72)^6^. However, therapeutic responses remain incomplete and variable among patients. Over 100 additional drug candidates targeting FXR, PPARs, incretin pathways, and other mechanisms are currently in phase II/III trials, though clinical attrition rates remain high^7^. Together, these observations underscore a persistent need to identify causal genes and pathways to enable mechanism-based patient stratification and therapeutic development.

The heritability for MASLD estimates from twin and family studies ranges from 20% to 70%^8^, yet MASLD GWAS have lagged behind those for other metabolic traits because robust phenotyping still requires liver biopsy or advanced imaging. The largest multi-ancestry GWAS meta-analysis has uncovered only 17 loci, explaining < 20% of heritability^9^. Proxy-phenotype GWAS using chronic alanine aminotransferase (ALT), fasting triglycerides (TG), or fatty liver index (FLI) as surrogates for MASLD have led to the identification of additional loci, expanding the catalog significantly^10–12^, although many causal genes remain unresolved.

Post-GWAS analytics help bridge this gap. Mendelian randomization (MR) and transcriptome-wide association studies (TWAS) couple GWAS statistics with molecular QTLs to test gene-trait causality. For example, MR screening of 1263 druggable genes pinpointed *ACE2* and *IFNAR2* as causal mediators of COVID-19 hospitalization, prioritizing repurposing opportunities for early management^13^. A tissue-specific TWAS integrating brain eQTLs identified *NK2R* as a driver of adiposity and glucose dysregulation, inspiring long-acting *NK2R* agonists that produced metabolic benefits *in vivo*^14^. These examples illustrate how post-GWAS analyses can translate association signals into actionable biology.

In this study, we integrated liver transcriptomic meta-analyses across MASLD stages and Summary-data-based Mendelian Randomization (SMR)/TWAS across MASLD and 13 related phenotypes, nominating 39 putative regulatory genes. We then prioritized candidates using a multi-evidence framework incorporating expression-phenotype correlation meta-analysis, PWAS support, and external genetic evidence (GWAS Catalog and ExPheWAS), and leveraged single-nucleus RNA-seq together with Genebridge to guide experimental selection. *MLIP* was prioritized for follow-up experimental validations based on its hepatocyte-enriched expression and convergent support across analyses. *MLIP* was induced in a lipid overload hepatocyte model, and its knockdown reduced lipid droplet accumulation and downregulated lipid metabolic programs, including PPAR signaling, triglyceride metabolism, and lipid transport/localization, highlighting *MLIP* as a regulator of hepatocellular lipid metabolism.

## Results

### Meta-Analysis of Differentially Expressed Genes in MASLD Progression

Considering the heterogeneity of MASLD and the biological variability among populations across different datasets, analyses based on a limited number of datasets may introduce potential biases. To identify genes that exhibit conserved expression changes in MASLD, we systematically retrieved all available MASLD-related transcriptomic datasets from public databases. After quality control, a total of 29 datasets comprising 2640 samples were included in the analysis. Dataset details are provided in Figure 1A and Table S1. Next, we reclassified the collected samples into three groups: Healthy Control, MASL, and MASH. The Healthy Control group included individuals classified as normal control in the original studies or those with a nonalcoholic fatty liver disease activity score (NAS) of 0. The MASL group comprised samples originally categorized as NAFL or MASL, as well as those with a NAS between 1 and 4. The MASH group included samples classified as NASH or MASH in the original studies or those with a NAS between 5 and 8. After classifying all MASLD samples, we performed three pairwise differential expression analyses within each dataset: (i) MASL vs. Healthy Control, representing gene expression changes associated with early disease progression; (ii) MASH vs. MASL, capturing gene expression changes during late-stage disease progression; and (iii) MASH vs. Healthy Control, reflecting gene expression alterations across the entire disease course. For each contrast, study-specific log2 transformed fold change (log2FC) estimates were synthesized with a restricted-maximum-likelihood (REML) random effects model; the pooled effect size is denoted randomSummary, and its meta-analytic significance is reported as random-P. Following meta-analysis, we kept genes tested in over 50% of the datasets with absolute randomSummary values over 0.2 and FDR below 0.05. This analysis identified 391 early MASLD progression differentially expressed genes (DEGs) (258 upregulated, 133 downregulated), 697 all MASLD stage DEGs (469 upregulated, 228 downregulated), and 315 late MASLD progression DEGs (241 upregulated, 74 downregulated) (Figure 1B). The full meta-analysis results for all genes across the three MASLD contrasts are available in Table S3. Consistent with well-established MASLD studies, our pipeline re-captured several well-established MASLD markers or regulators, such as *FABP*, *IL32* and *SPP1*^15–17^, confirming the accuracy of our pooled effect-size estimates. In addition, we highlighted multiple less-characterized dysregulated genes, expanding the set of candidate drivers for downstream analyses (Figure 1C). We also asked which genes are associated with MASLD-related fibrotic progression by performing fibrosis stage–based differential-expression meta-analyses (F1/2 vs F0, F3/4 vs F1/2, and F3/4 vs F0). The complete fibrosis meta-analysis results are provided in Table S4, and an overview of DEG counts and representative signatures is shown in Supplementary Figure 1.

**Figure 1.**
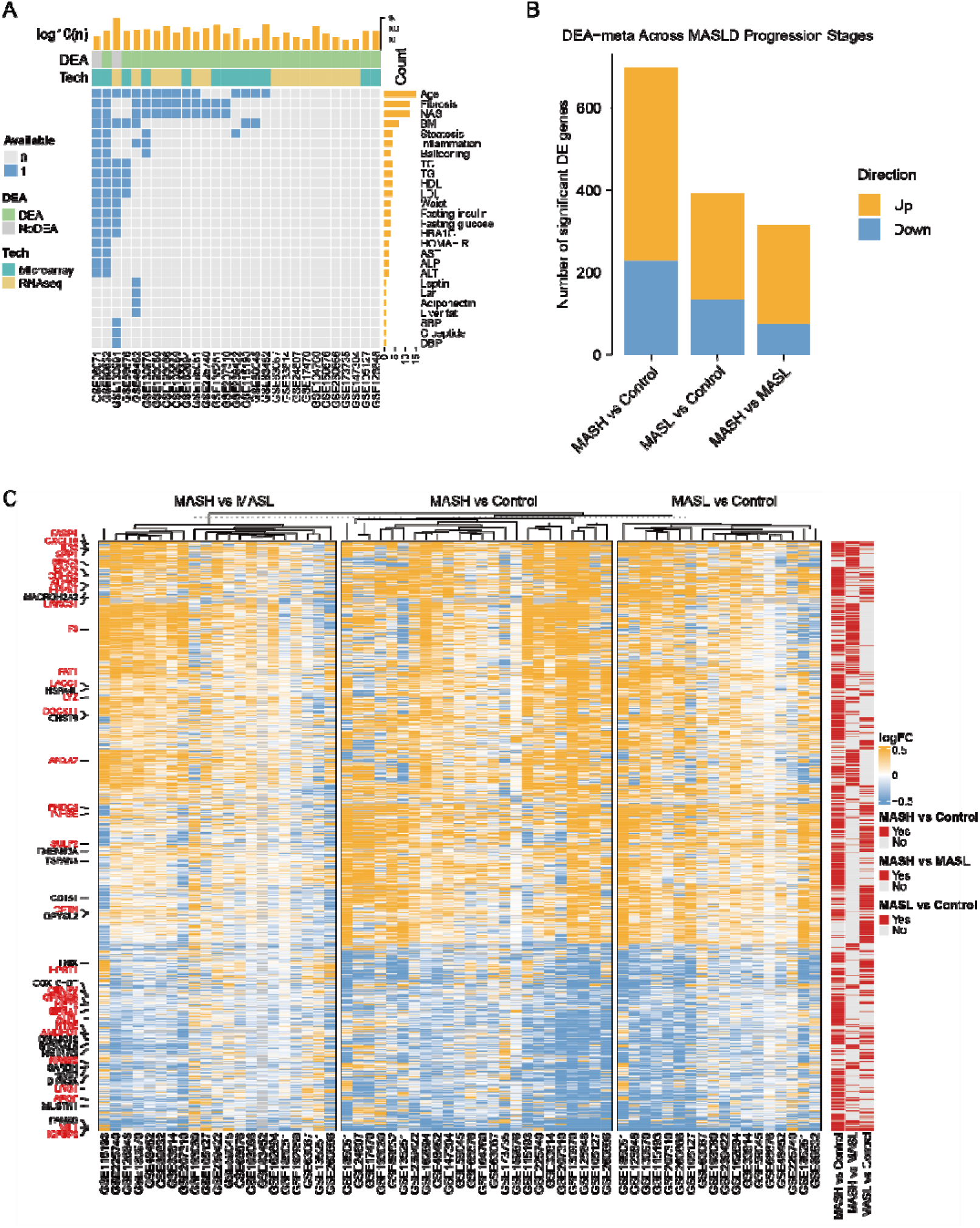
Transcriptomic Meta-Analysis Identifies Dysregulated Genes in MASLD Progression. (A) Overview of datasets included in transcriptomic meta-analysis. Columns represent individual datasets and rows correspond to available clinical parameters; blue indicates the presence and grey the absence of a given parameter in each dataset. The top bar chart shows the sample size of each dataset (log10(n)). The second annotation row denotes whether the dataset was included in differential expression analysis (green = yes; grey = no). The third annotation row indicates the profiling platform (green = microarray; yellow = RNA-seq). (B) Bar plots show the number of significantly up-regulated (yellow) and down-regulated (blue) genes (FDR < 0.05, |randomSummary| ≥ 0.2) identified in each pairwise meta-analysis comparison. (C) Heatmap displaying all significantly differentially expressed genes for each pairwise comparison (MASH vs. Healthy Control, MASL vs. Healthy Control, and MASH vs. MASL); left-side row annotations highlight the top 10 up-regulated (yellow) and top 10 down-regulated (blue) genes in each comparison. Genes with prior evidence for MASLD association (curated from published literature) are highlighted in red. The three annotation columns on the right indicate whether each gene is significant in the corresponding comparison (red = significant; grey = not significant).

To distinguish stage-specific pathways from shared transcriptional programs along the MASLD trajectory, we contrasted the three meta-analytic DEG sets. Venn diagrams of up- and downregulated genes (Figure 2A, B) indicated that many meta-DEGs were specific to either the early transition (MASL vs control) or the steatohepatitis transition (MASH vs MASL), whereas a smaller subset was shared across all three contrasts. We operationally defined stage-specific genes as those significant in only one transition contrast under identical thresholds, and subjected these sets to pathway enrichment analysis (Table S5). Upregulated genes specific to early-stage MASLD were enriched for SREBF-driven lipogenic programs, cholesterol biosynthesis, and extracellular-matrix organization, whereas MASH-specific upregulated genes preferentially mapped to inflammatory and extracellular matrix–remodeling pathways (Figure 2C). Conversely, early-stage-specific downregulated genes were enriched for circadian clock components and FOXO-related transcriptional regulation, while MASH-specific downregulated genes were linked to xenobiotic metabolism, amino-acid metabolism and gluconeogenesis (Figure 2D). Finally, we defined a 27-gene core progression signature comprising genes that were significant and directionally concordant across all three pairwise meta-analyses. This signature—including *AKR1B10*, *FABP4*, *FCAMR*, *FMO1*, *IL32*, and *SPP1*—showed largely monotonic shifts from controls to early-stage MASLD and further to MASH across independent cohorts (Figure 2E). Together, these results define a conserved transcriptional signature of MASLD progression and nominate candidate biomarkers and therapeutic entry points for future study.

**Figure 2.**
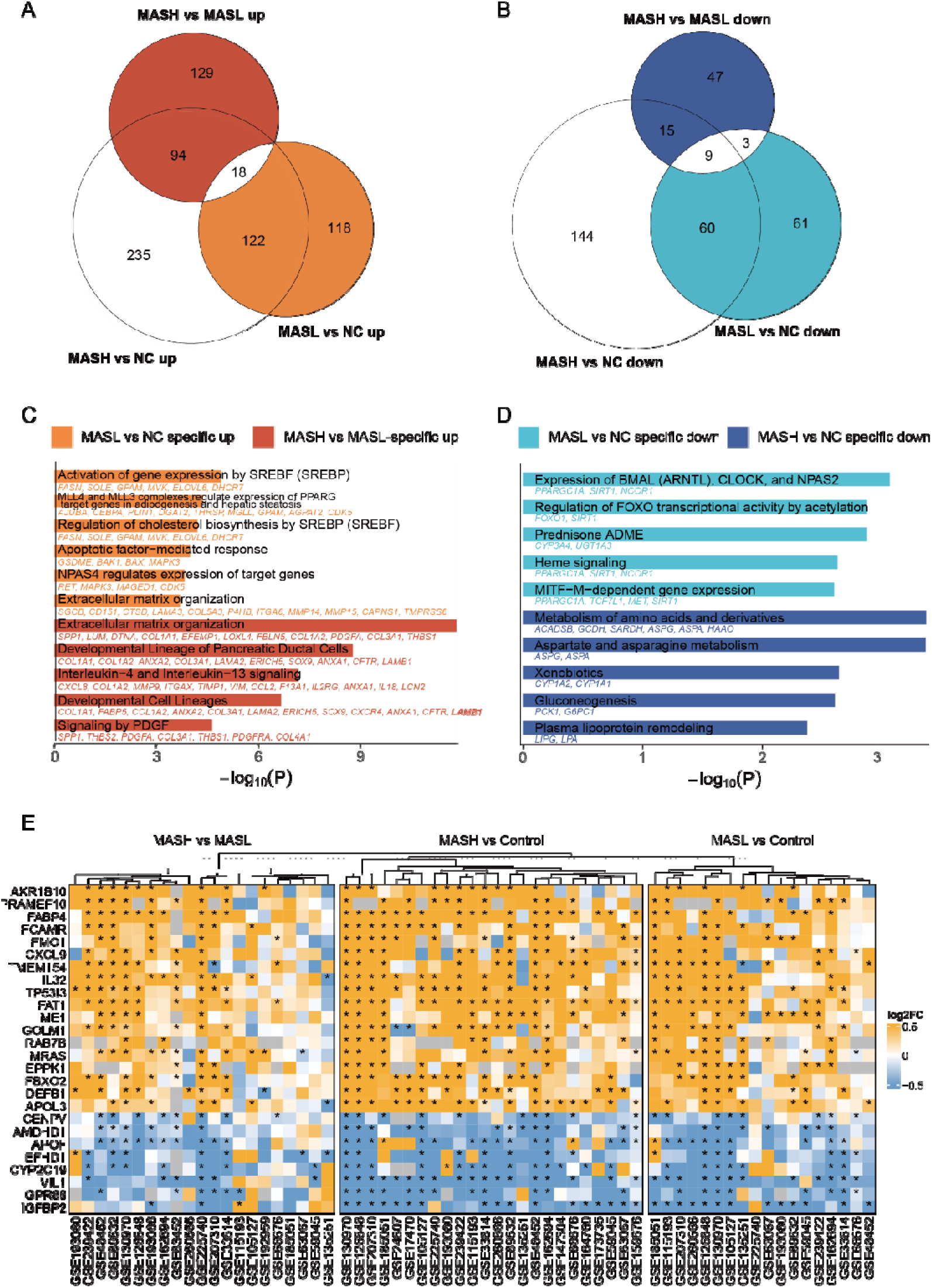
Stage-specific and core transcriptional programs during MASLD progression. (A-B) Venn diagrams showing the overlap of significantly up-regulated (A) and down-regulated (B) DEGs identified by meta-analysis of MASL vs NC, MASH vs MASL, and MASH vs NC comparisons. (C-D) Reactome pathway enrichment for early stage—specific and late stage—specific DEGs, showing up-regulated (C) and down-regulated (D) pathways. Bars represent −log10(p values) for selected pathways, with member genes listed next to each bar. (E) Heatmap of a 27-gene conserved MASLD progression signature. rows represent genes and columns datasets (grouped by contrast), with asterisks indicating statistical significance.

Beyond the ORA of stage-specific DEG sets, we next asked whether stage-specific versus shared transcriptional pathways would remain evident when considering the entire ranked transcriptome. For each contrast, genes were ranked by the randomsummary from the REML random effects meta-analysis and subjected to GSEA, enabling a comprehensive view of pathways engaged at disease initiation (MASL vs Control), those emerging during progression to steatohepatitis (MASH vs MASL), and a conserved set shared across stages (Figure 3A, B, Table S6). A subset of pathways was concordantly upregulated across all three contrasts, including DNA-damage and stress-response programs, senescence-associated secretory phenotype (SASP), extracellular-matrix/collagen organization, and inflammatory signaling modules (Figure 3A, C). In contrast, MASL-specific upregulated pathways were dominated by mitochondrial fatty-acid beta-oxidation and related lipid-metabolic processes, whereas MASH-specific upregulated pathways were enriched for TNF receptor, inflammasome, and cytokine signaling, consistent with an inflammatory and fibrogenic switch at the MASH transition. Downregulated pathways showed a distinct stage dependence: in MASL, suppression was enriched for circadian clock and hormone-related signaling (including leptin pathways), whereas in MASH repression shifted toward carbohydrate, lipid, and amino-acid metabolic pathways, consistent with progressive loss of hepatocellular metabolic capacity (Figure 3B, C). Notably, several pathways (e.g., mitochondrial fatty-acid beta-oxidation and gluconeogenesis) displayed biphasic behavior—upregulated in MASL vs Control but downregulated in MASH vs MASL—suggesting an initial compensatory activation of energy-generating programs that ultimately fails as disease advances (Figure 3C, Figure S2). Finally, NES heatmaps across cohorts (Figure 3C) revealed broadly consistent enrichment patterns within pathway groups, supporting recurrent, stage-dependent pathway modules across independent datasets.

**Figure 3.**
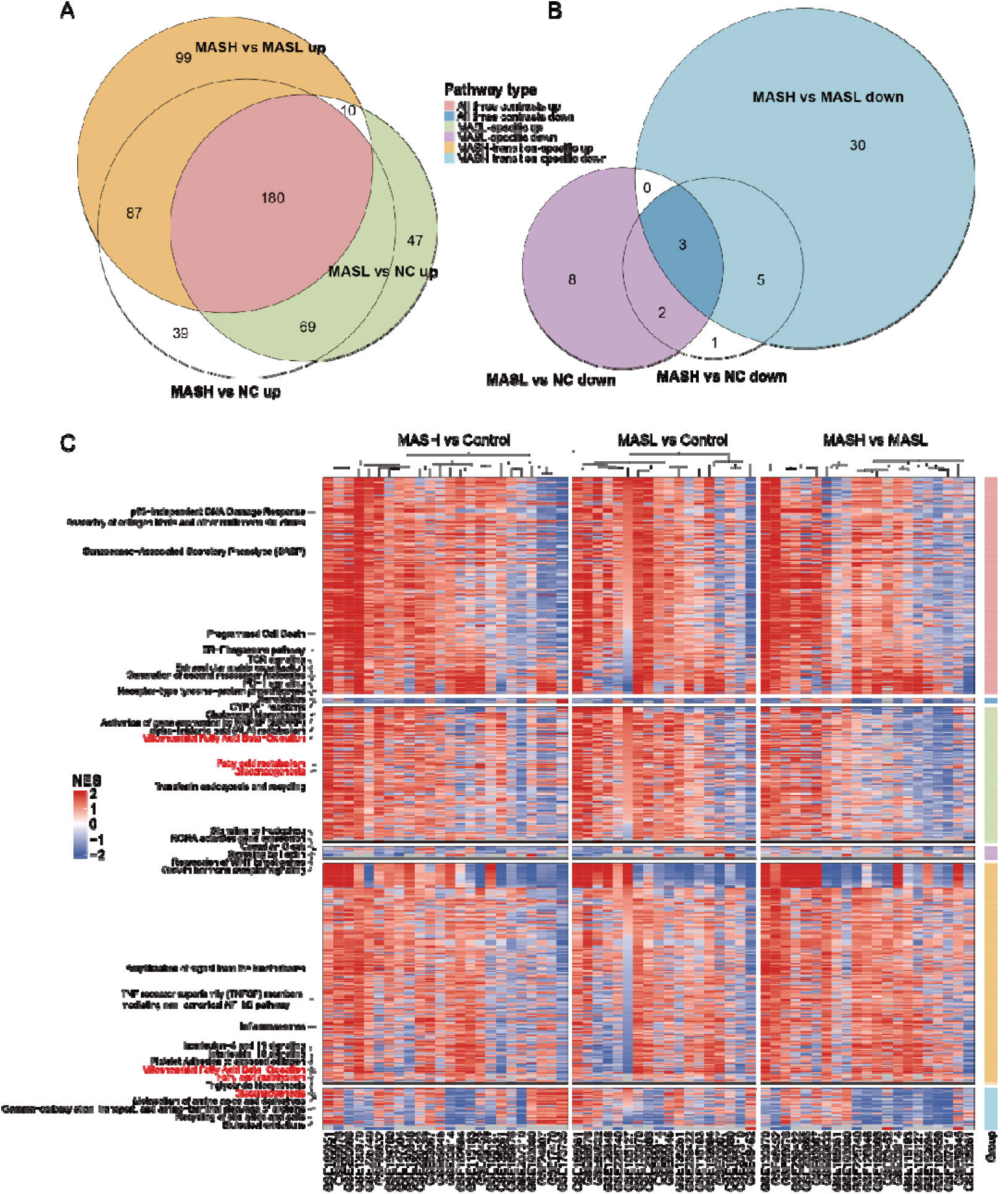
Stage-specific and shared pathway programs along the MASLD trajectory. (A-B) Venn diagrams showing the overlap of significantly enriched Reactome pathways across the three meta-analytic contrasts (MASL vs Control, MASH vs MASL, and MASH vs Control). Panel A shows pathways with positive NES (upregulated), and panel B shows pathways with negative NES (downregulated). Colors indicate pathway categories (shared across contrasts or preferentially enriched in a single contrast). (C) Heatmap of NES for representative Reactome pathways from each category in (A–B) across all individual datasets. Columns denote studies, grouped by the three contrasts, and rows denote pathways clustered into MASL-specific, MASH-transition-specific, and shared modules; the right-side color bar indicates the pathway category. Red and blue represent positive and negative NES, respectively. Pathways highlighted in red text indicate biphasic enrichment (up in MASL vs Control but down in MASH vs MASL).

### Integrative correlation meta-analysis links hepatic gene expression to clinical features of MASLD

To quantify how clinical features of MASLD are reflected in the hepatic transcriptome, we correlated gene expression with 19 clinical parameters spanning anthropometrics (BMI, waist circumference), metabolic traits (fasting glucose, fasting insulin, HbA1c, HOMA-IR, triglycerides, total cholesterol, HDL, LDL), histological scores (steatosis, lobular inflammation, ballooning, fibrosis stage, NAS), and biochemical markers of liver injury (ALT, AST), from 18 human cohorts. Within each cohort, we computed Spearman correlation coefficients for each gene-trait pair and combined study-specific estimates using a REML random-effects model to obtain meta-analyzed correlations and corresponding confidence intervals.

Using NAS as an example of composite disease activity, Figure 4A shows the cohort-level correlations between hepatic *FABP4* expression and NAS across 12 independent MASLD datasets (left), together with a meta-analytic summary and the strongest NAS-associated genes ranked by meta-analyzed correlation (right). *FABP4* showed a consistently positive association with NAS across cohorts and ranked among the top positively correlated genes, together with extracellular-matrix genes (e.g., *COL1A1*, *COL1A2*, *COL4A1*), whereas metabolic genes including *APOF* and *GNMT* were among the most negatively correlated ones.

**Figure 4.**
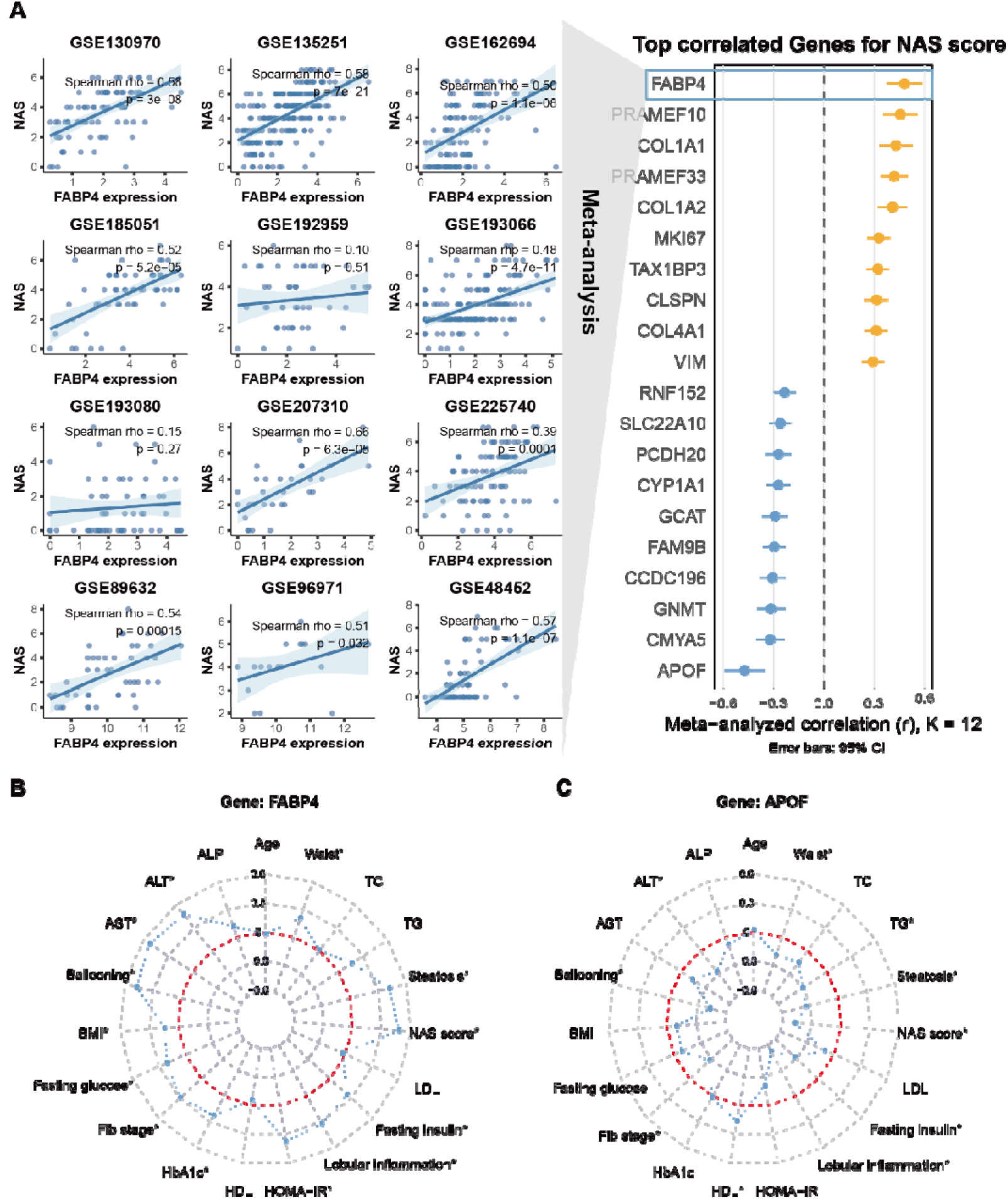
Correlation meta-analysis links hepatic gene expression to NAS and broader clinical profiles. (A) Left, cohort-level Spearman correlations between hepatic *FABP4* expression and NAS across 12 independent MASLD datasets; each panel shows the correlation coefficient and p value, with a trend line for visualization. Right, meta-analyzed correlations for the top NAS-associated genes. Points indicate meta-analyzed Spearman’s rho with 95% confidence intervals; orange and blue denote positive and negative correlations, respectively. (B-C) Radar plots summarizing meta-analyzed correlations between *FABP4* (B) or *APOF* (C) and 19 clinical parameters. Radius denotes the meta-analyzed correlation coefficient; asterisks mark FDR-significant associations.

To summarize broader clinical association profiles, we visualized meta-analyzed correlations across all 19 traits using radar plots. *FABP4* exhibited positive correlations with histological activity and fibrosis measures as well as metabolic risk traits (Figure 4B). In contrast, *APOF* showed predominantly negative correlations with NAS and related histological and metabolic traits (Figure 4C), consistent with its reduced expression in samples with greater metabolic and histological burden. Meta-analyzed gene–trait correlation results for all genes and all clinical parameters are provided in Table S7 (see also Supplementary Figure S3 for top genes per trait). We further extended this framework to the pathway level by correlating ssGSEA-derived Reactome pathway activity scores with each clinical trait; the complete pathway–trait correlation results are provided in Table S8. All gene-and pathway-level correlation profiles are additionally searchable through the interactive web portal (https://masldportal.net/).

### Integrated TWAS, SMR, and transcriptomic meta-analysis converge on 39 candidate MASLD regulators

To prioritize candidate genes with putative causal relevance to MASLD despite the limited power of biopsy/imaging-defined MASLD GWAS, we implemented an integrative strategy that leverages genetically related phenotypes as complementary discovery channels. Prior proxy-phenotype GWAS using ALT, triglycerides, or fatty liver-related indices have replicated established loci and identified additional associations, supporting the utility of proxy-informed approaches^10–12,18^. Building on this rationale, we sought to increase discovery power while simultaneously assessing pleiotropy across MASLD-relevant disease dimensions.

We curated 13 MASLD-related phenotypes from prior genetic correlation, Mendelian randomization, and clinical association studies^19–21^, including liver injury markers (ALT, AST, GGT), lipid traits (TG, HDL-C, LDL-C), glycemic control (HbA1c), systemic factors (CRP, SHBG), and MASLD complications (CAD, cirrhosis, T2D, MetS). Using GTEx liver eQTLs as a shared anchor, we performed SMR with HEIDI testing and TWAS for MASLD and each related phenotype (Figure 5A-B; Table S9–S10). Under our analysis settings, MASLD GWAS alone yielded no significant SMR/TWAS genes, consistent with the limited power of current biopsy/imaging-defined MASLD GWAS. We therefore integrated genetic results across phenotypes and intersected SMR/TWAS-supported genes with the tri-stage hepatic differential-expression meta-analysis, yielding a refined set of 39 candidates supported across genetic and transcriptomic layers (Figure 5C).

**Figure 5.**
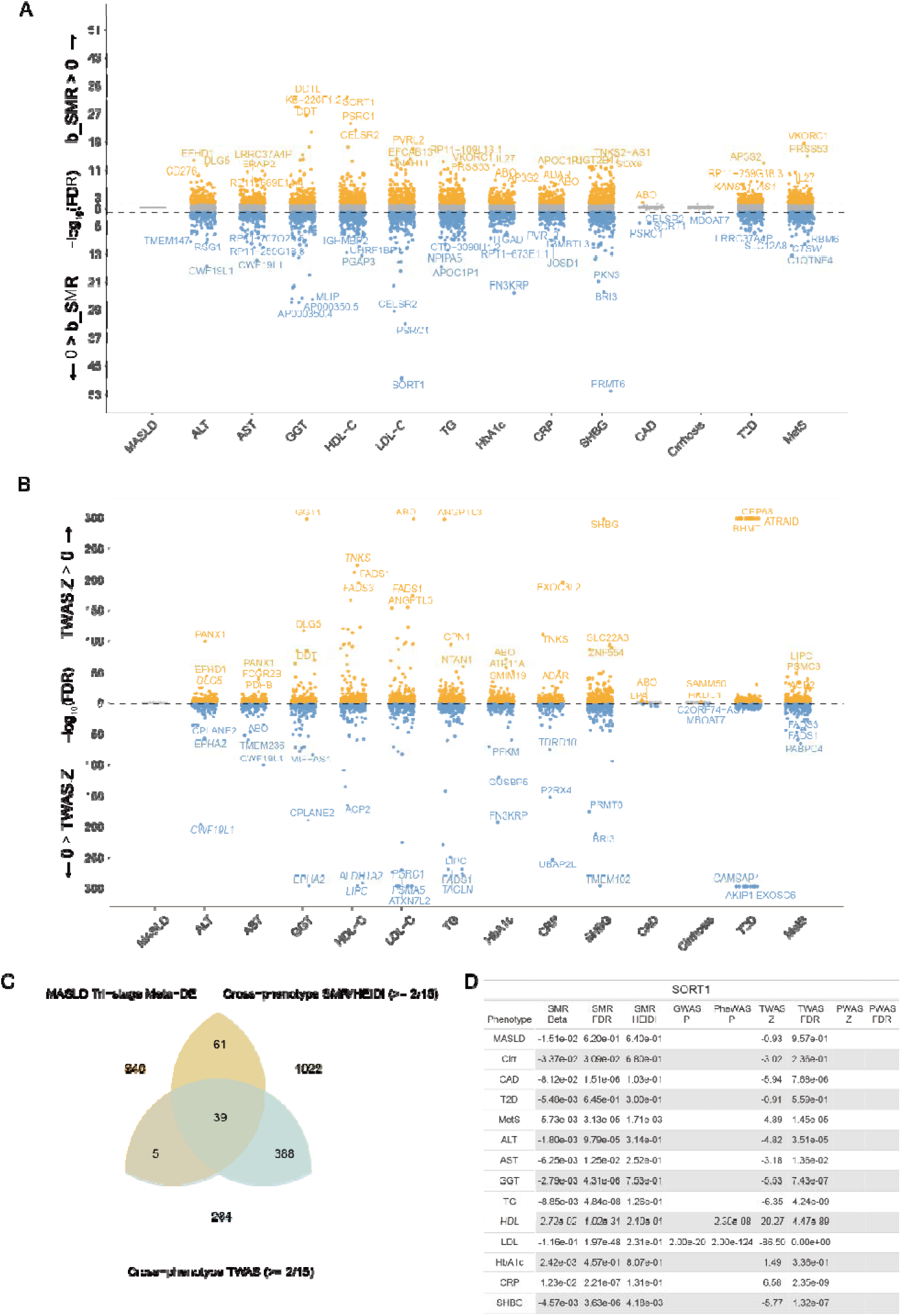
Integrated TWAS, SMR, and transcriptomic meta-analysis converge on 39 candidate MASLD regulators. (A–B) Bidirectional Manhattan plots summarizing the SMR (A) and TWAS (B) results for MASLD and 13 MASLD-related phenotypes using GTEx-liver eQTL data. Yellow points indicate positively associated genes and blue points indicate negatively associated genes; the top three positive and negative signals are labelled for each phenotype. (C) Venn diagram showing the intersection of the SMR and TWAS findings (restricted to genes significant in ≥ 2 of the 14 phenotypes) with the tri-stage MASLD differential expression meta-analysis, yielding a 39-gene candidate set. (D) *SORT1* is shown as an illustrative example from this set. The bar chart summarizes its direction of association across multiple phenotypes in SMR and TWAS, while external ExPheWAS and GWAS Catalog evidence highlights its broad pleiotropy across lipid traits, CAD and MASLD-related traits.

To illustrate multi-layer support, we highlighted *SORT1* as an example. Genetic and experimental evidence indicates that *SORT1* is a hepatic regulator of lipoprotein trafficking and VLDL secretion, and that higher hepatic *SORT1* expression is associated with a more favorable lipid profile (lower LDL-C and VLDL/TG-related traits) and reduced CAD risk^22^. Consistent with this direction, our liver eQTL—anchored SMR/TWAS analyses showed that genetically predicted hepatic *SORT1* expression was inversely associated with CAD and atherogenic lipid traits. Separately, transcriptome-based correlation meta-analysis linked hepatic *SORT1* expression to multiple MASLD-related clinical measures, supporting a broader phenotypic footprint for *SORT1* across metabolic and liver disease domains. Several additional genes in the 39-candidate gene set (e.g., *EFHD1*, *GAS6*, *GNMT*) have prior functional support in MASLD, reinforcing the biological relevance of our integrated prioritization.

### Multi-layer evidence integration prioritizes 39 candidate genes

To refine the 39 candidates emerging from genetic and transcriptomic analyses, we constructed a composite prioritization score integrating six complementary evidence layers: (i) tri-stage MASLD differential-expression meta-analysis, (ii) gene-clinical parameter correlation meta-analysis, (iii) cross-phenotype liver eQTL-anchored SMR with HEIDI testing, (iv) cross-phenotype TWAS, (v) cross-phenotype PWAS (Figure S4; Table S11), and (vi) external genetic evidence from the GWAS Catalog (Table S12) and ExPheWAS (Table S13). For each layer, genes were assigned a binary support indicator using pre-specified criteria: differential expression required meta-FDR < 0.05 and |randomSummary| ≥ 0.2; correlation required meta-FDR < 0.05 in ≥2 clinical traits; SMR required FDR < 0.05 with HEIDI P > 0.05 in ≥2 phenotypes; TWAS and PWAS required FDR < 0.05 in ≥2 phenotypes; GWAS Catalog support required *P* < 5×10^−8^ with the gene annotated as a “mapped gene” in ≥2 traits; and ExPheWAS support required FDR < 0.05 in ≥2 traits. Indicators were summed to yield the final priority score used to rank candidates (Table 1). *EFHD1* ranked first, reflecting consistent support across evidence layers. In line with this prioritization, an independent human MASLD single-cell eQTL study reported a hepatocyte maladaptation–associated loss of an *EFHD1* eQTL with FOXO1-linked genotype-specific regulation^23^, further supporting *EFHD1* as a MASLD susceptibility candidate.

**Table 1.**
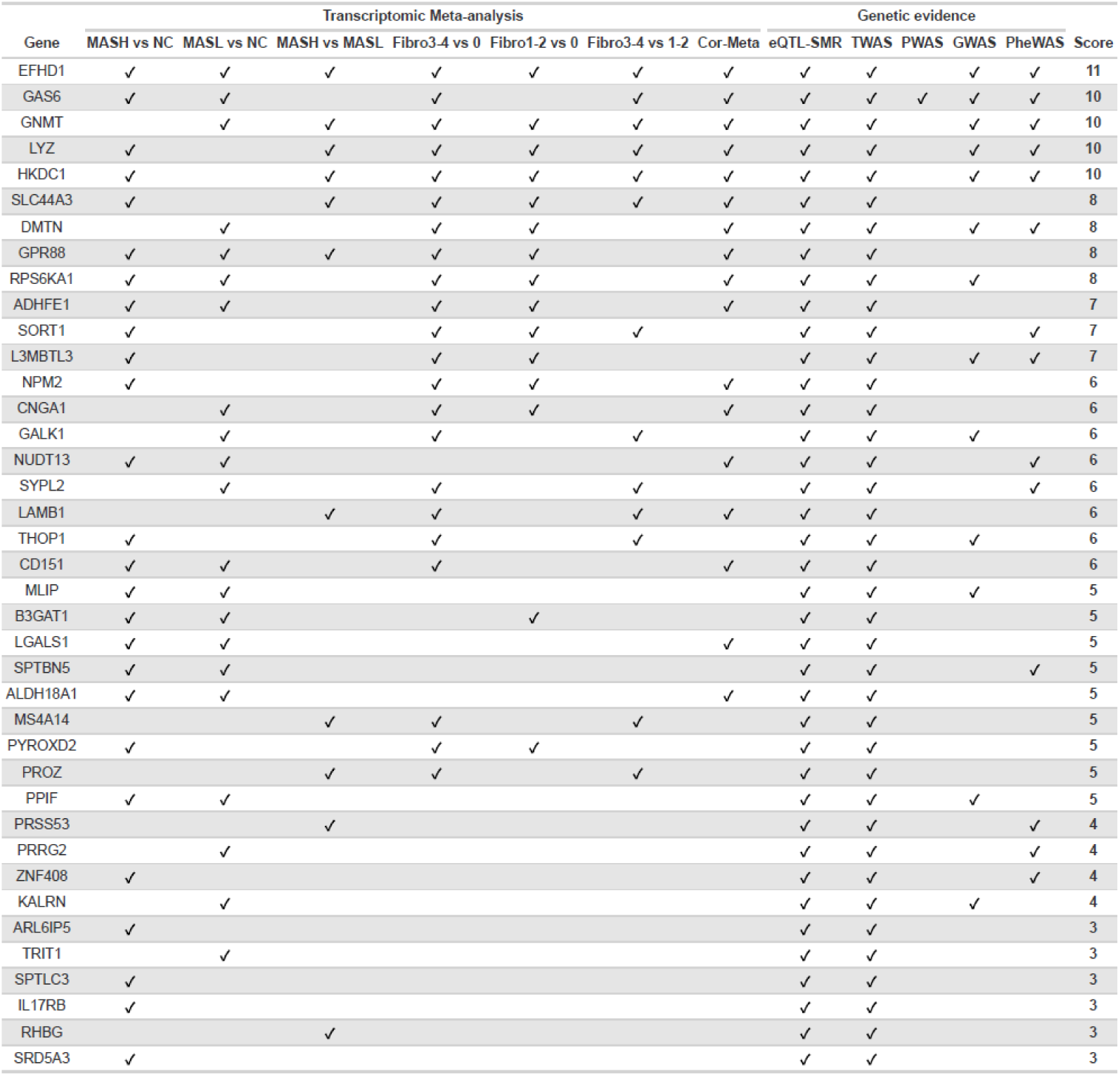
Integrative evidence matrix for 39 candidate MASLD regulator genes. Rows are genes; columns on the left summarize transcriptomic meta-analysis support (MASH vs. Control, MASL vs. Control, MASH vs. MASL, fibrosis stage contrasts, and correlation meta-analysis), and columns on the right capture genetic-level evidence (SMR, TWAS, PWAS, GWAS hits, and PheWAS signals). A checkmark indicates the gene met the predefined significance criteria in that specific analysis (see Methods). The “Score” is the sum of positive lines of evidence used in the prioritization framework, with genes ordered by descending score.

### Single-nucleus RNA-seq mapping of 39 prioritized MASLD candidate genes across liver cell types and disease stages

To place the 39 prioritized candidate genes in a cellular context, we analyzed a human MASLD liver snRNA-seq dataset spanning controls and multiple disease stages, including MASH with fibrosis or cirrhosis. UMAP embeddings illustrate the annotated liver cell types and disease-stage composition (Figure S5A-B). Mapping candidate-gene expression across cell types revealed marked heterogeneity and clear cell-type enrichment patterns (Figure S5C). For example, *GNMT* and *ADHFE1* were preferentially expressed in hepatocytes, consistent with hepatocellular metabolic functions, whereas *LYZ* was enriched in Kupffer cells, suggesting an immune-associated function.

### Integrated multi omics resource and public portal

To enable transparent access to all analysis layers and support community reuse beyond the 39 prioritized candidates, we built an interactive MASLD Portal, available at https://masldportal.net. The portal provides gene-centric queries and summarizes, for any gene, three complementary evidence views: (i) forest plots from differential-expression meta-analyses across MASLD contrasts (MASH vs Control, MASL vs Control, and MASH vs MASL) and fibrosis-stage contrasts (F1/2 vs F0, F3/4 vs F1/2, and F3/4 vs F0); (ii) radar plots capturing the direction and magnitude of meta-analyzed gene-clinical trait correlations; and (iii) an integrated evidence table compiling SMR/TWAS signals, ARIC plasma PWAS support, GWAS Catalog annotations, and ExPheWAS annotation. Supplementary Fig. S6 shows the portal interface and available analysis modules.

This resource recapitulates established MASLD regulators while also highlighting candidates with limited functional characterization. For example, querying *MLIP* returns its stage- and fibrosis-associated expression patterns, multi-trait correlation profile, and cross-phenotype genetic/proteomic support in a single view, enabling rapid inspection of consistency, directionality, and breadth across data modalities (Figure 6).

**Figure 6.**
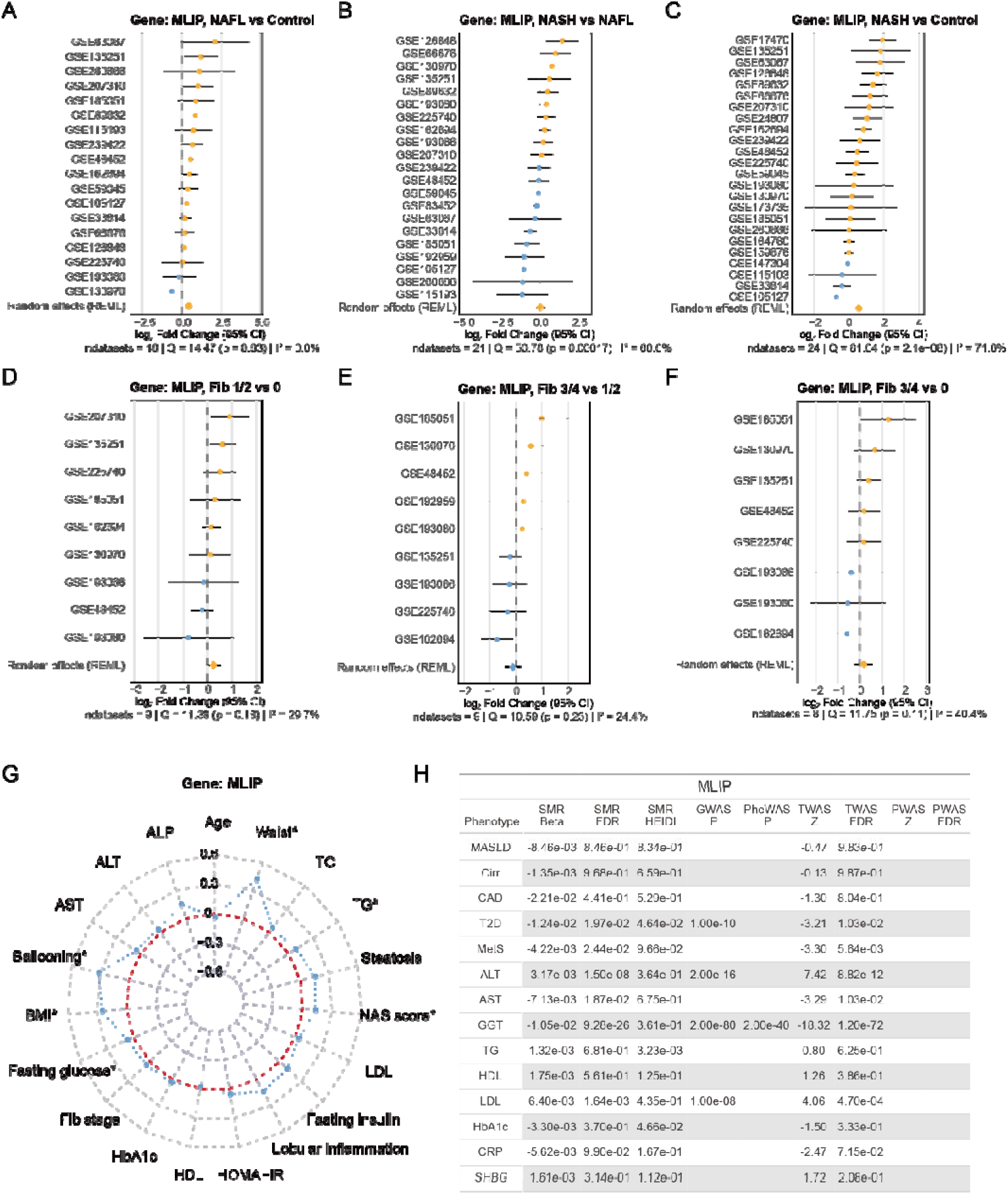
Gene-centric integrated evidence dashboard of *MLIP*. For any queried gene, example shown for *MLIP*, the panel includes: (A-F) Forest plots of differential expression meta-analysis across MASLD disease stages (MASH, MASL and Control) and fibrosis progression (F0, F1/2, F3/4), with effect sizes (randomSummary) and 95% CI; (G) radar plots summarizing correlation meta-analysis with clinical parameters; and (H) a consolidated table of multi omics support, including significance in SMR/TWAS, PWAS, GWAS Catalog annotations and PheWAS associations.

### MLIP modulates hepatocellular lipid accumulation and lipid metabolic programs

Among the 39 prioritized candidates, we prioritized *MLIP* for functional follow-up based on convergent evidence implicating a hepatocyte-centered, early-stage signal. In the human MASLD snRNA-seq dataset, *MLIP* showed predominantly hepatocyte-enriched expression, and within hepatocytes its expression was significantly higher in MASLD F0 than in healthy controls, with lower levels in cirrhotic stages (Figure 7A; Figure S5C). Consistent with this cellular context, GeneBridge analysis in human liver linked *MLIP* to lipid metabolism—related modules, including fatty-acid oxidation and related lipid metabolic processes^24^ (Figure 7B). Together, these observations motivated mechanistic evaluation of *MLIP* in hepatocyte models.

**Figure 7.**
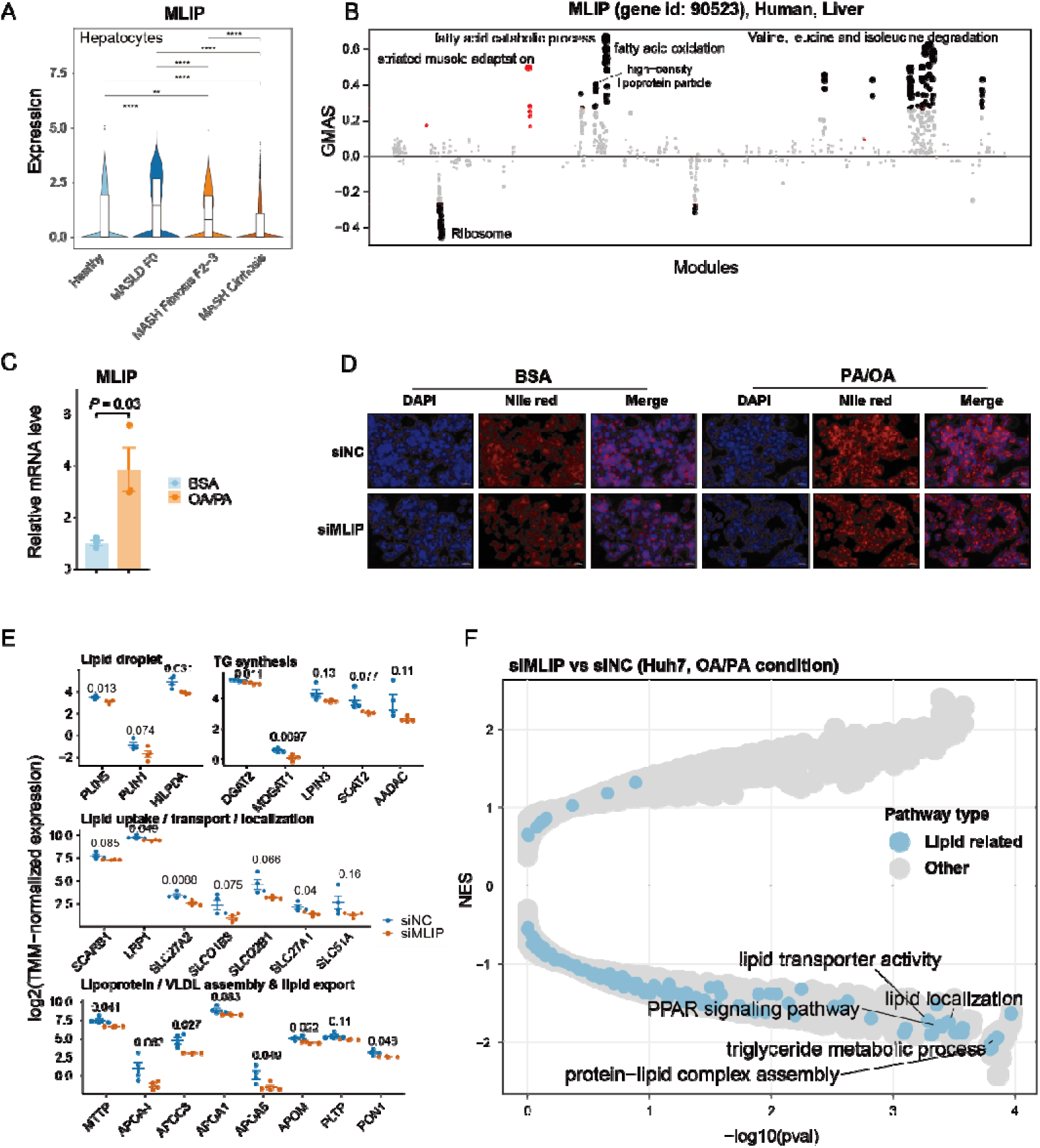
*MLIP* modulates hepatocellular lipid accumulation and lipid metabolic programs. (A) *MLIP* expression in hepatocytes across disease stages in a human MASLD snRNA-seq dataset. Statistical significance is indicated as *P < 0.05, **P < 0.01, ***P < 0.001, ****P < 0.0001. (B) GeneBridge module associations for *MLIP* in human liver. The y axis denotes the gene-module association score (GMAS), GMAS > 0.268 indicate positive association and GMAS < −0.268 indicate negative association. (C) qPCR quantification of *MLIP* mRNA in Huh7 cells after OA/PA treatment relative to control. (D) Representative fluorescence images of Huh7 cells stained with Nile Red (lipid droplets) and DAPI (nuclei) under vehicle control (BSA) or OA/PA conditions with siNC or siMLIP; merged images are shown. (E) Expression of representative genes involved in lipid droplet dynamics, triglyceride synthesis, lipid transport/localization, and lipoprotein/VLDL assembly/export in OA/PA-treated Huh7 cells comparing siMLIP versus siNC. (F) GSEA comparing siMLIP versus siNC under OA/PA. Each dot represents a pathway; lipid-related pathways are highlighted, with selected terms labeled.

To model lipid overload *in vitro*, we treated HuH7 cells with a mixture of oleic acid (OA) and palmitic acid (PA). *MLIP* expression increased upon OA/PA exposure (Figure 7C). Using Nile Red staining, we observed robust lipid droplet accumulation under OA/PA conditions, which was attenuated by *MLIP* knockdown (Figure 7D). In parallel, targeted transcriptional readouts showed coordinated changes in lipid-handling gene programs upon *MLIP* knockdown, including reduced expression of genes involved in lipid droplet dynamics and triglyceride synthesis, as well as altered expression of genes related to lipid uptake/transport/localization and lipoprotein/VLDL assembly and export (Figure 7E). At the pathway level, RNA-seq followed by enrichment analysis further indicated suppression of lipid-related pathways in siMLIP versus siNC under OA/PA, including PPAR signaling, triglyceride metabolic processes, and lipid transport/localization modules (Figure 7F). Collectively, these data place *MLIP* within hepatocellular lipid metabolic programs and support its functional involvement in lipid accumulation under lipotoxic conditions.

## Discussion

In this study, we developed a liver expression—anchored multi-omic framework to prioritize candidate regulators of MASLD by integrating differential expression meta-analysis, gene expression–clinical trait correlation meta-analysis, liver eQTL–anchored genetic evidence (SMR with HEIDI and TWAS), PWAS, and external annotations (GWAS Catalog and ExPheWAS). This layered strategy converged on 39 candidates supported across multiple data types, including established regulators (e.g., *SORT1*, *EFHD1*) and less-characterized genes such as *MLIP*. We further mapped these candidates in a human MASLD snRNA-seq atlas to define their dominant cellular contexts and stage-associated expression patterns. Finally, we performed mechanistic follow-up in hepatocyte lipid-overload models and show that *MLIP* knockdown attenuates lipid droplet accumulation and dampens lipid metabolic programs, supporting a functional role for *MLIP* in hepatocellular lipid handling.

At the transcriptomic level, stage- and fibrosis-centered meta-analyses revealed coordinated dysregulation consistent with distinct pathological programs across the MASLD continuum. Early-stage comparisons were dominated by lipid metabolic remodeling, whereas progression toward MASH and advanced fibrosis increasingly reflected inflammatory activation and extracellular matrix organization. Gene—clinical trait correlation meta-analysis added clinical resolution by positioning individual genes along complementary pathogenic axes: correlations with fibrosis stage nominate genes linked to matrix-associated processes; associations with ALT/AST suggest relationships with hepatocellular injury; and links to dyslipidemia or insulin resistance point to metabolic stress. Together, these trait-aligned profiles help interpret clinical heterogeneity and highlight that MASLD progression reflects intertwined metabolic, inflammatory, and stromal programs rather than a single linear pathway.

Integrating liver eQTL—anchored SMR and TWAS across MASLD and a curated set of MASLD-related phenotypes expanded gene discovery beyond the limited power of biopsy/imaging-defined MASLD GWAS alone. Proxy-phenotype GWAS (e.g., ALT, triglycerides, fatty liver indices) have demonstrated that phenotype expansion can recover known loci and uncover additional signals^10,11,18^; here, anchoring inference on hepatic gene expression enabled us to evaluate cross-phenotype support while keeping the analysis grounded in liver regulatory mechanisms. *SORT1* illustrates this multi-layer logic: prior genetic and experimental studies link hepatic *SORT1* to lipoprotein metabolism and cardiometabolic risk, and in our analyses genetically predicted hepatic *SORT1* expression showed protective-direction associations with CAD and atherogenic lipid traits. Transcriptome-based correlations further connected *SORT1* expression to MASLD-related clinical and histological measures, highlighting a broad phenotypic footprint consistent with pleiotropic involvement across metabolic and liver disease domains.

More broadly, this expression-anchored, cross-phenotype integration provides a principled way to prioritize genes based on consistency and directionality across multiple evidence layers while retaining liver regulatory relevance. Because such multi-layer support extends beyond the 39 highlighted candidates, we provide an interactive portal to enable transparent exploration and independent follow-up of the complete evidence landscape.

The gene-centric public portal (masldportal.net) extends the utility of this work by making the full multi-layer evidence landscape readily accessible for any gene of interest. By integrating differential-expression meta-analysis, clinical correlation profiles, liver eQTL–anchored genetic support, PWAS signals, and external annotations into a single interface, the portal enables rapid inspection of evidence directionality, consistency across datasets, and cross-phenotype support for any gene of interest, facilitating transparent hypothesis generation and independent validation.

This study has several strengths. First, the framework integrates complementary layers—cross-cohort transcriptomic meta-analysis, and liver eQTL—anchored SMR/TWAS—thereby prioritizing candidates supported across independent analytical axes rather than a single dataset or modality. Second, REML random-effects models increase robustness to between-study heterogeneity, while pre-specified multi-layer criteria reduce reliance on any single signal. Third, single-nucleus mapping provided cellular context for candidate interpretation and helped distinguish baseline cell-type enrichment from disease-associated expression shifts. Finally, incorporating external resources and releasing an interactive portal improves transparency and reusability, enabling the community to interrogate the full evidence space.

Limitations should be noted. First, cohort and technical heterogeneity—including variable phenotyping, biopsy scoring practices, and platform differences—may introduce residual confounding despite harmonization and random-effects modeling. Second, SMR/TWAS infer expression-mediated genetic associations under specific assumptions and can be affected by linkage disequilibrium structure or horizontal pleiotropy; thus, not all prioritized genes can be interpreted as causal effector. Third, clinical parameters were not uniformly available across cohorts, which may limit correlation resolution for some traits. Finally, although we provide functional evidence for *MLIP* in hepatocyte lipid-overload models, systematic *in vivo* and cell-type–specific validation across the candidate set remains necessary. In addition, our operational use of MASL/MASH classifications reflects available cohort annotations and scoring thresholds, which may not perfectly align with all contemporary nomenclature frameworks.

Future work should pursue deeper mechanistic dissection of top candidates using cell-type–resolved perturbation (e.g., overexpression and *in vivo* validation) and integrate spatial and multi-modal omics to capture microenvironmental contributions and post-transcriptional regulation. Expanding liver eQTL resources and incorporating diverse ancestries and prospective cohorts will further strengthen genetic prioritization and generalizability. Finally, refined multi-gene panels derived from the prioritized set should be evaluated in independent cohorts for staging, prognosis, and treatment-response stratification, with iterative updates deployed through the web portal.

In summary, we integrate cross-cohort transcriptomic meta-analysis, liver expression–anchored genetic inference, clinical trait mapping, proteomic associations, and targeted functional follow-up into a unified framework that prioritizes 39 candidate MASLD regulators. By providing both cellular context and a public gene-centric portal, this work offers a reusable resource for mechanistic discovery and supports translation toward biomarker development and therapeutic target nomination.

## Methods

### GWAS summary statistics of MASLD and related traits

We collected publicly available, European-ancestry GWAS summary statistics for MASLD and a set of related phenotypes selected based on prior evidence of genetic correlation or Mendelian randomization–supported causal relationships with MASLD. The phenotype panel comprised liver injury markers (ALT, AST, GGT), lipid traits (triglycerides, LDL-C, HDL-C—with underlying variant/imputation framework anchored to the 1000 Genomes Project Phase 3 reference), glycemic traits (HbA1c), inflammation/hormonal markers (CRP, SHBG), and comorbidities (coronary artery disease, cirrhosis, type 2 diabetes, and metabolic syndrome). All datasets and their provenance (including source consortia, sample sizes, and phenotype definitions) are detailed in Table S2.

### Meta-analysis of Differential Gene Expression

For identify preserved differentially expressed genes, three-steps methods performed in this work, i) Data acquisition and filtering, publicly available human liver RNAseq and microarray datasets were queried from GEO using the search terms: “MASLD/metabolic dysfunction-associated fatty liver disease”, “fatty liver/steatosis”, “NASH/non-alcoholic steatohepatitis”, “MAFLD/metabolic dysfunction-associated fatty liver disease”, and “Homo sapiens.” Datasets which lacking matched control samples or ambiguous disease staging information were excluded. And only data derived from whole liver tissues were retained, while datasets generated from *in vitro* cell lines or sorted cell populations were discarded to avoid confounding effects. Samples showing significant deviation in principal component analysis (PCA) were flagged as technical or biological outliers and subsequently removed. The retrieved datasets and detail information are summarized in Table S1. ii) Differential gene expression analysis, to standardize the classification of disease progression across datasets, we categorized the samples into three stages: Healthy Control, Early Stage, and Late Stage. Healthy Control was defined based on the original study’s classification as either Normal Control (NC) or having a NAFLD Activity Score (NAS) of 0. Early stage included samples originally classified as NAFL/MAFL/MASL or those with a NAS score ranging from 1 to 4 and late stage encompassed samples identified as NASH/MASH in the original studies or those with a NAS score between 5 and 8. This classification approach ensured consistency in disease staging while aligning with widely accepted histopathological criteria for MASLD progression. To ensure analytical robustness, for RNA-seq data, low-expressed genes were filtered, retaining those with more than 5 reads in at least half of samples. Gene expression data from RNA-seq datasets were normalized using the trimmed mean of M-values (TMM) method via the calcNormFactors function in edgeR (v3.34.1)^25^. For microarray datasets, we used log2-transformed expression values from pre-processed data provided by the original sources.

Differential expression analysis for both TMM-normalized RNA-seq datasets and processed microarray datasets was performed using the limma package (v3.38.3)^26^. Meta-analysis of differential expression was performed using a restricted maximum likelihood (REML) random-effects model to combine study-specific log2 fold change (log2FC) estimates^27,28^. For each pairwise contrast, the log2FC and its standard error from each dataset were input, and the pooled effect size from the REML model is reported as randomSummary, with the corresponding meta-analytic significance denoted random-P. To focus on reproducible signals, only genes observed in at least 50% of the contributing datasets for a given comparison were considered. Multiple testing correction was applied to the random-P values using the Benjamini–Hochberg procedure to obtain FDR. Genes with |randomSummary| ≥ 0.2 and FDR < 0.05 were deemed significantly differentially expressed.

### Correlation meta-analysis of hepatic gene expression with clinical features

Within each MASLD cohort, we quantified associations between hepatic gene expression and 19 clinical parameters using Spearman’s correlation method for each gene–trait pair. Cohort-specific correlation estimates were then combined across studies using a random-effects meta-analysis fitted by REML to obtain meta-analyzed correlations with 95% confidence intervals and P values. Multiple testing was controlled using Benjamini–Hochberg FDR. In parallel, we derived Reactome pathway activity scores within each cohort by single-sample gene set enrichment analysis (ssGSEA) implemented in the GSVA R package (ssGSEA method, v1.50.5)^29^, and applied the same Spearman correlation and REML random-effects meta-analysis workflow to test pathway-trait associations.

#### Gene Set Enrichment analysis (GSEA)

Enrichment analysis was conducted using the fgsea package (version 1.30.0)^30^ on DEGs between each comparison. All DEGs of each comparison were preranked by log2FC or randomSummary and analyzed with genesets from the GO, KEGG, and REACTOME databases.

### Reactome pathway enrichment analysis

Reactome pathway enrichment analysis was performed on the consensus MASLD signatures (genes consistently upregulated or downregulated across MASH vs Control, MASH vs MASL, and MASL vs Control) derived from meta-analysis results using the clusterProfiler R package (version: 4.10.1)^31^.

### Single-nucleus RNA sequencing analysis

Public single-nucleus RNA sequencing (snRNA-seq) datasets were downloaded from GEO (GSE256398). In short, 6 healthy control samples and 11 MASLD samples, including 3 MASLD F0 samples, and 4 MASH Fibrosis F2-F3 samples, and 4 MASH Cirrhosis samples were included in this dataset^32^. After obtaining the raw feature-barcode matrices file for each sample, we constructed Seurat objects for each sample using the Seurat R package (version 4.4.0)^33^. Quality control was applied to remove low-quality cells. Specifically, cells were excluded if they had fewer than 200 or more than 6,500 detected genes (nFeature_RNA), more than 40,000 total UMI counts (nCount_RNA), or if over 20% of their genes originated from mitochondrial genes (percent.mt). Next, 3000 highly variable genes (HVGs) were extracted using the FindVariableFeatures function. Based on the expression of these HVGs, dimensionality reduction was performed using principal component analysis (PCA) and we kept 30 principal components (PCs) for further analysis. The RunUMAP function and the FindClusters function (resolution = 0.25) were used to cluster single cells. Applying well-known marker genesto annotate cell clusters into known cell types^34^.

### Transcriptome-wide Mendelian randomization

SMR was used to integrate GWAS and eQTL data for causal inference of MASLD-related traits^35^. We focused on cis-eQTLs associated with common SNPs (minor allele frequency > 0.1%) to prioritize genes with likely regulatory effects. Expression data for liver tissues from the GTEx v8 database. HEIDI colocalization testing was applied to identify shared causal variants between eQTL and GWAS signals, with an FDR < 0.05 threshold for significance and a p-value > 0.05 for HEIDI to exclude non-colocalized signals.

### TWAS and PWAS

TWAS was performed with the FUSION^36^, using GTEx-liver cis-eQTL weights to impute genetically regulated gene expression and test its association with MASLD and related traits by integrating the corresponding GWAS summary statistics. Associations passing FDR < 0.05 were considered significant. We then carried the same framework forward to a PWAS by substituting FUSION pQTL weights derived from ARIC plasma proteomics, thereby extending the analysis to protein-level associations^37^.

### Composite gene prioritization score

To rank the 39 candidates, we compiled a fine-grained evidence matrix comprising transcriptomic meta-analysis contrasts and genetic evidence items (Table 1; Tables S3, S4, S7, S11–S13). For each gene and each evidence item, we assigned a binary indicator (1/0) using pre-specified criteria. Transcriptomic evidence items included pairwise MASLD-stage differential-expression meta-analyses (MASL vs NC, MASH vs NC, and MASH vs MASL) and fibrosis-stage differential-expression meta-analyses (Fib1-2 vs 0, Fib3-4 vs 0, and Fib3-4 vs 1-2), and gene-clinical trait correlation meta-analysis. Genetic evidence items included SMR with HEIDI testing (eQTL-SMR), TWAS, PWAS, GWAS Catalog mapping, and ExPheWAS. Differential-expression items were considered supported if meta-FDR < 0.05 and |randomSummary| ≥ 0.2 for the corresponding contrast; Cor-Meta support required meta-FDR < 0.05 for the corresponding clinical trait set; SMR support required FDR < 0.05 with HEIDI P > 0.05; TWAS/PWAS support required FDR < 0.05; GWAS Catalog support required P < 5×10^−8^ with the gene annotated as a mapped gene; and ExPheWAS support required FDR < 0.05. The final priority score was computed as the sum of all supported evidence items across columns.

### GeneBridge

Gene-pathway associations were evaluated using the Gene-Module Association Determination (G-MAD) analysis implemented in GeneBridge^24^. We used the human liver resource in GeneBridge, which integrates 51 liver transcriptomic datasets (each with >80 samples) to provide meta-analyzed gene-module association scores (GMAS) for curated GO, KEGG, and Reactome gene sets. Associations were interpreted using the recommended thresholds: GMAS > 0.268 for significant positive association and GMAS < −0.268 for significant negative association.

### RNAseq

Cells were harvested and total RNA was extracted by using RNAmini kit (Qiagen, Germany) and sequenced on Illumina Novaseq 6000 platform to generate paired-end 2×150 bp reads. Reads were quality-filtered using fastp (v0.22.0)^38^ and aligned to the human reference genome (GRCh38) using STAR (v2.7.10b)^39^. Gene-level counts were quantified using featureCounts (Subread v2.0.6). Differential expression analysis was performed with edgeR (v3.34.1) using raw counts.

### Cell Culture, Transfection and Treatments

Huh7 cells were obtained from the American Type Culture Collection (ATCC). Cells were maintained at 37 °C with 5% CO_2_ in Dulbecco’s modified Eagle medium (DMEM; Corning, NY, USA) supplemented with 10% fetal bovine serum (FBS; Serana Europe GmbH, Brandenburg, Germany) and 1% penicillin–streptomycin (Sangon Biotech, Shanghai, China). For gene knockdown, cells were transfected with siRNAs targeting *MLIP* (siMLIP) or a non-targeting negative control siRNA (siNC) using siRNA-mate plus (GenePharma, Shanghai, China) according to the manufacturer’s protocol. Briefly, 30 pmol siRNA was complexed with 3 μL siRNA-mate plus per well (final ratio 30 pmol:3 μL) and added to cells. The siMLIP sense and antisense sequences were 5’-GAGUCCAGAAACUGUAAAUTT-3’ and 5’-AUUUACAGUUUCUGGACUCTT-3’, respectively. The negative control siRNA sense and antisense sequences were 5’-UUCUCCGAACGUGUCACGUTT-3’ and 5’-ACGUGACACGUUCGGAGAATT-3’, respectively. At 24 h post-transfection, cells were treated for 48 h with a palmitic acid/oleic acid (PA/OA) mixture to induce lipid accumulation (PA, 333 μM; Sigma-Aldrich, Taufkirchen, Germany; OA, 666 μM; Beyotime, Shanghai, China). Bovine serum albumin (BSA; MP Biomedicals, Santa Ana, CA, USA) was used as the vehicle control. Cells were subsequently harvested for RT-qPCR and transcriptomic analysis.

### Cellular Nile Red staining

For lipid droplet visualization, cells were washed with phosphate-buffered saline (PBS), fixed with 4% paraformaldehyde for 10 min at room temperature, and washed three times with PBS. Cells were stained with Nile Red (1 μg/mL; MedChemExpress, Shanghai, China) for 10 min protected from light. Nuclei were counterstained with DAPI (Beyotime, Shanghai, China) for 5 min (final concentration per manufacturer’s instructions). After three washes with PBS, images were acquired using an inverted fluorescence microscope (Leica MATEO FL RUO, Shanghai, China) under identical acquisition settings across conditions. Representative DAPI, Nile Red, and merged images are shown.

### Real-time Quantitative PCR

Total RNA was isolated from Huh7 cells using AG RNAex Pro Reagent (Accurate Biology, Hunan, China) following the Trizol method. cDNA was synthesized using a reverse transcription kit (Accurate Biology, Hunan, China). Quantitative PCR was performed using SYBR Green chemistry (ABclonal, Wuhan, China) on a CFX Connect Real-Time PCR Detection System (Bio-Rad, Hercules, CA, USA). Relative gene expression was calculated using the 2^-ΔΔCT^ method and normalized to β-actin as the internal reference. Primer sequences are provided in Table S16.

### Statistical analysis

Statistical analyses and figure generation were performed using R (v4.3.1). Data are presented as mean ± s.e.m. Two-group comparisons were assessed by two-tailed Student’s t-test, multiple comparisons were corrected using Benjamini-Hochberg method. External gene–trait association evidence was retrieved from the GWAS Catalog (downloaded on 2025-06-28, Table S12) and ExPheWAS (accessed on 2025-06-25, Table S13) and incorporated into the composite prioritization framework as described above.

## Data availability

All transcriptomic datasets used for differential expression analysis, stage-resolved meta-analysis, and gene–clinical correlation meta-analysis were obtained from the Gene Expression Omnibus (GEO) database. Accession numbers and all GWAS summary-statistics sources curated for the present study, are listed in the Datasets section of the MASLD Portal (https://masldportal.net).

TWAS were performed using precomputed expression-weight reference models from FUSION based on GTEx v8. PWAS used protein prediction models derived from the ARIC cohort. External gene–trait association evidence was retrieved from the GWAS Catalog (downloaded on 2025-06-28) using the following trait identifiers: OBA_2050062, EFO_0004736, EFO_0001645, EFO_0001422, EFO_0004458, EFO_0004532, EFO_0004541, EFO_0004612, EFO_0004611, EFO_0003095, EFO_0000195, EFO_0004696, MONDO_0005148, and EFO_0004530. ExPheWAS gene–phenotype association data were downloaded from ExPheWAS (accessed on 2025-06-25).

## Authors’ contributions

A.Z. and H.L. conceived, designed, and supervised the project. Z.F., H.L., J.X. and Y.Z. collected and analyzed the data. F.C., A.D., J.D., and X.L. performed cell experiments. Z.F. and S.W. built and deployed the MASLDportal.net web portal. Z.F., F.C. and H.L. wrote the manuscript.

## Acknowledgement

The authors thank Jia Kang, and members of the Li lab for technical help and discussions.

---

**Figure S1.**
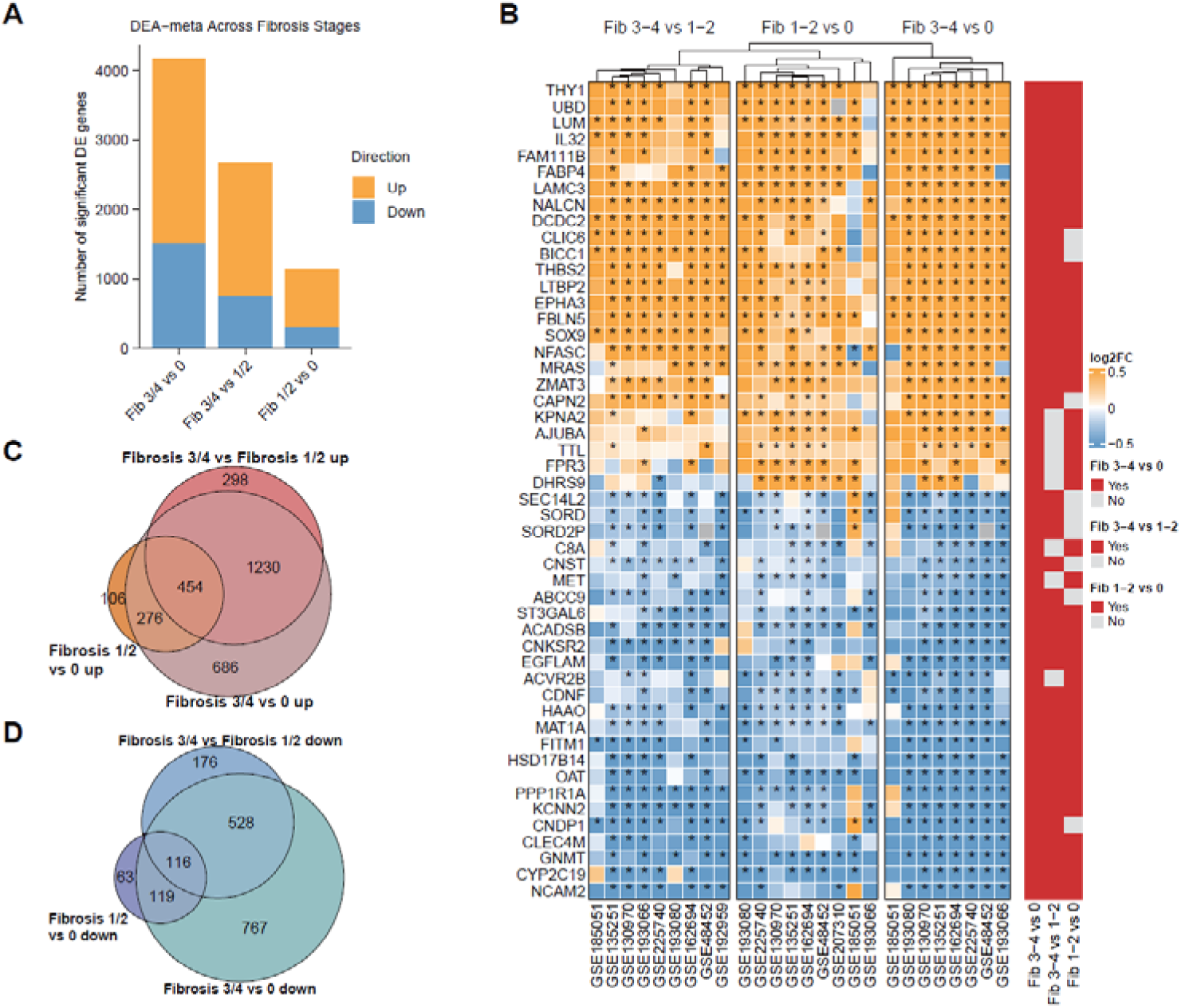
Differential expression meta-analysis across fibrosis stages in MASLD cohorts. (A) Numbers of significantly up- and down-regulated genes identified by random-effects (REML) meta-analysis for three fibrosis contrasts (F1-2 vs F0, F3-4 vs F1-2, and F3-4 vs F0). Significance was defined by FDR < 0.05 and |randomSummary| > 0.2. (B) Heatmap showing meta-analytic log2 fold changes for the top 10 up-regulated and top 10 down-regulated genes in each fibrosis contrast across individual datasets. Columns denote cohorts; rows denote genes. Asterisks indicate cohort-level differential expression significance (FDR < 0.05). (C-D) Venn diagrams depicting overlap of significantly up-regulated (C) and down-regulated (D) genes among the three fibrosis contrasts, highlighting shared and stage-transition-specific fibrosis-associated transcriptional programs.

**Figure S2.**
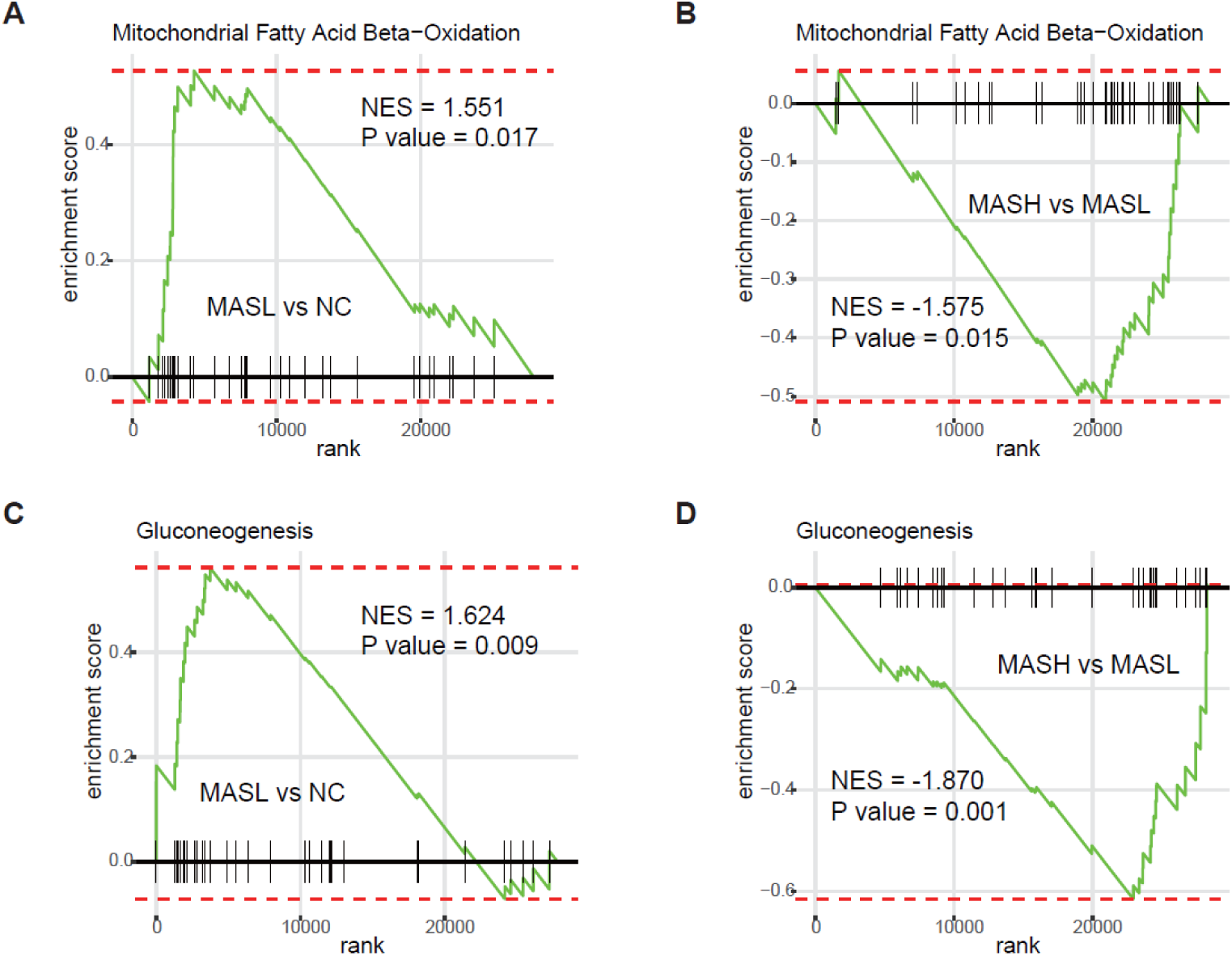
Biphasic pathway changes across the MASLD trajectory. Representative GSEA enrichment plots illustrating pathways that show opposite directions between early-stage MASLD and the transition to MASH. (A–B) Mitochondrial fatty acid β-oxidation is enriched in MASL vs control (A) but depleted in MASH vs MASL (B). (C–D) Gluconeogenesis is enriched in MASL vs control (C) but depleted in MASH vs MASL (D).

**Figure S3.**
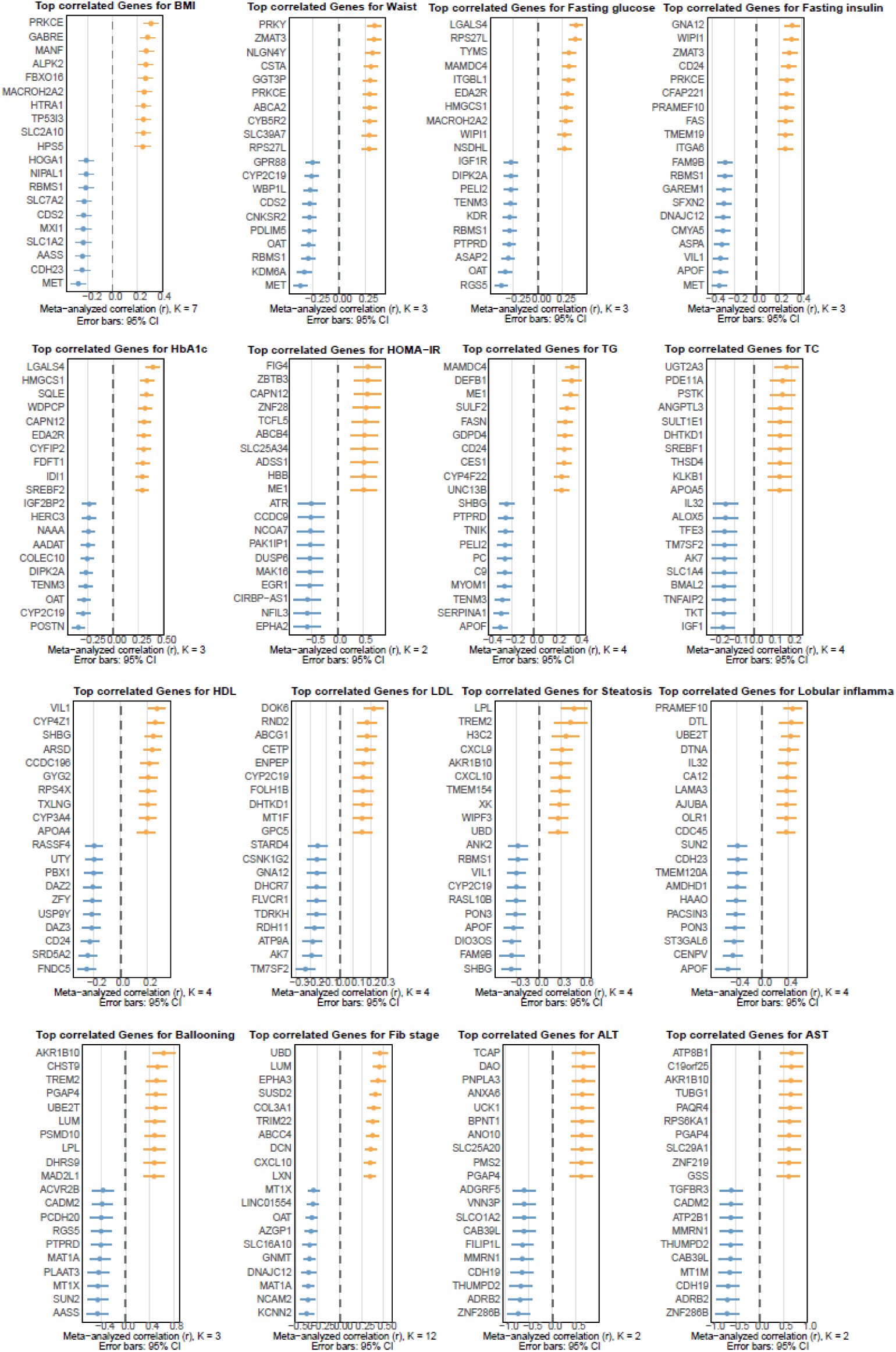
Top gene–clinical trait correlations across MASLD cohorts. Forest plots show the top positively (orange) and negatively (blue) correlated hepatic genes for each clinical parameter across cohorts. Points denote the meta-analyzed correlation (r) with 95% confidence intervals; K indicates the number of contributing datasets for each trait.

**Figure S4.**
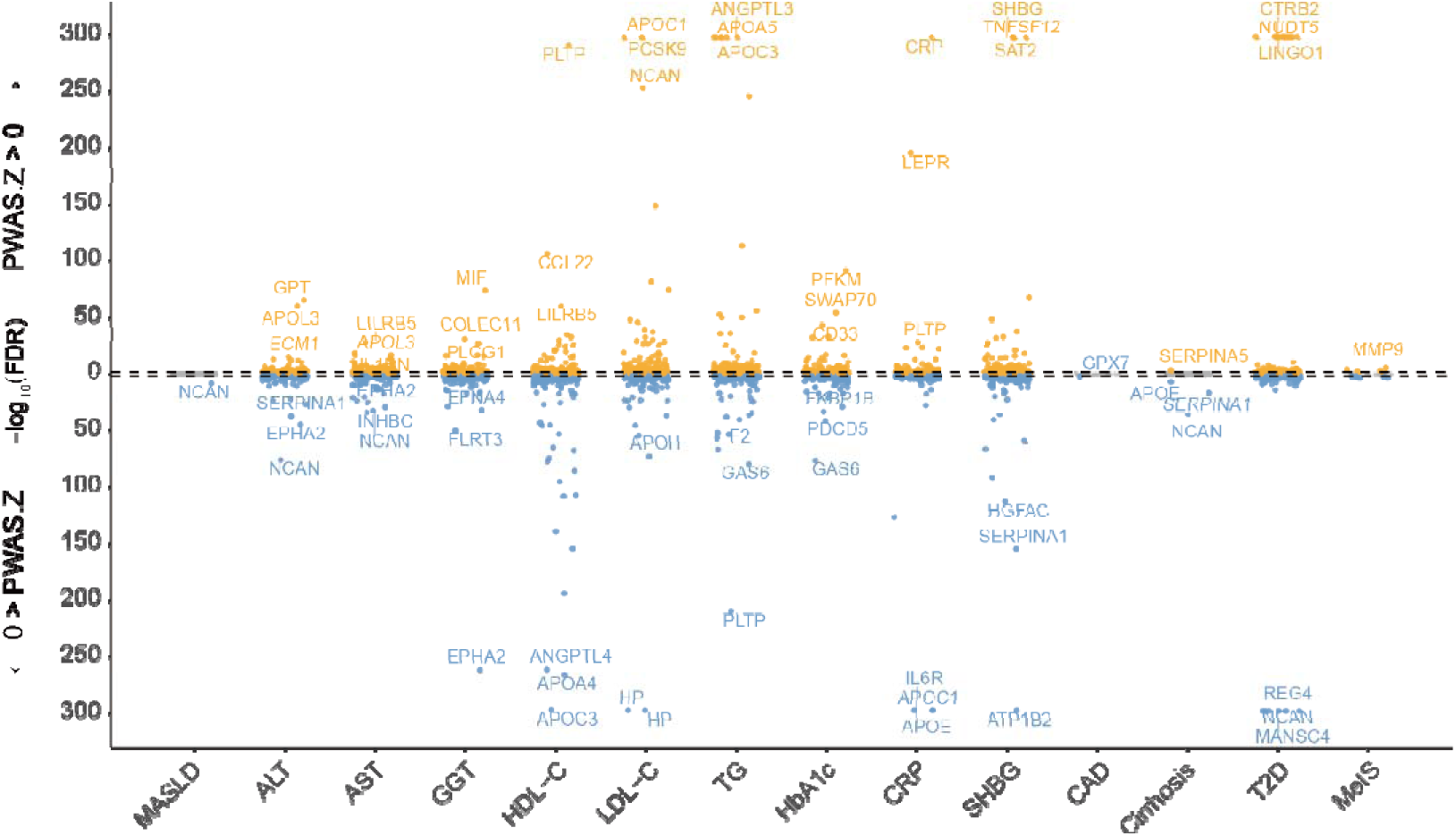
Plasma PWAS across MASLD and related phenotypes. Bidirectional Manhattan plots summarizing the PWAS results for MASLD and 13 MASLD-related phenotypes using ARIC proteomics data.

**Figure S5.**
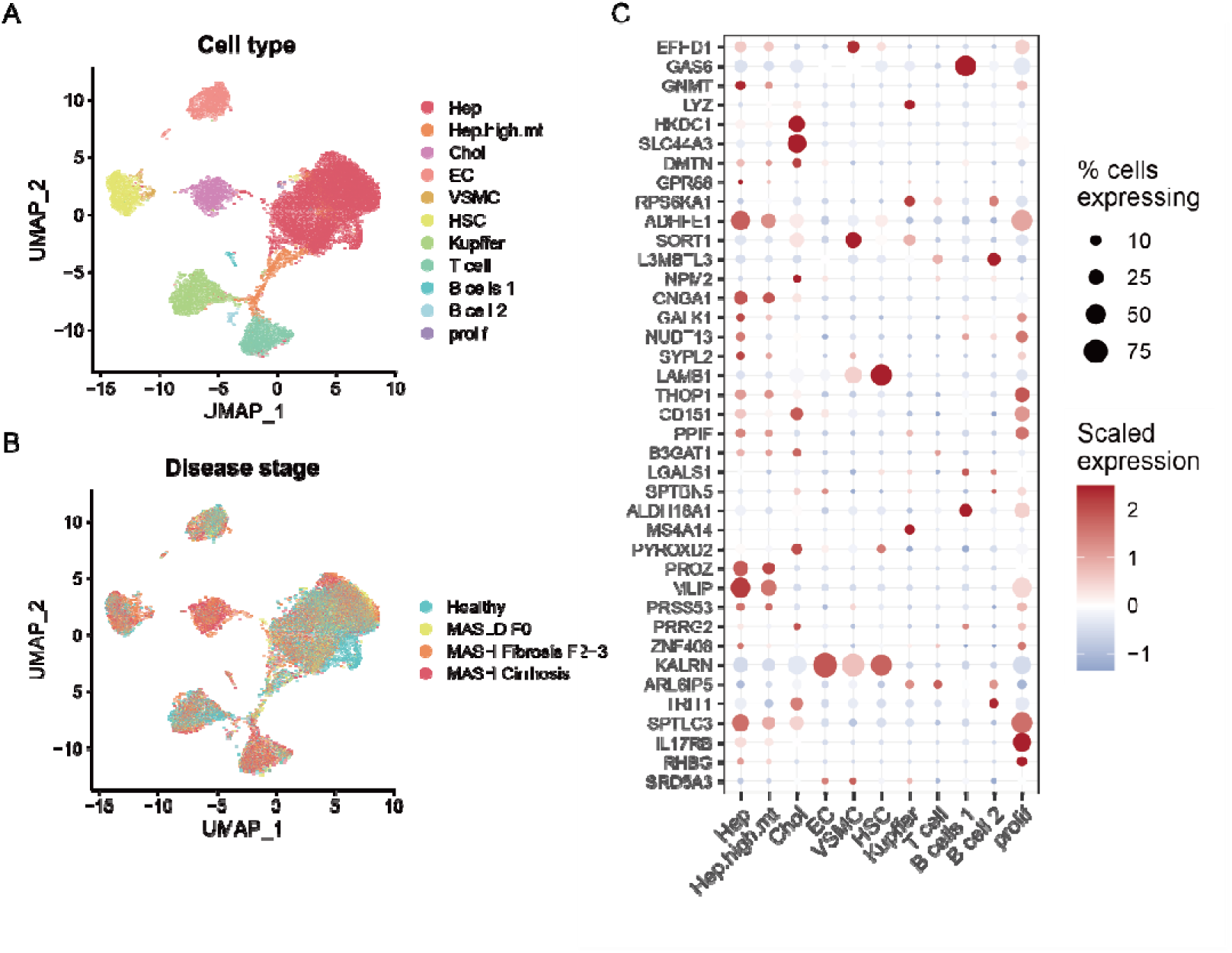
Single-nucleus RNA-seq mapping of 39 prioritized MASLD candidate genes across liver cell types and disease stages. (A-B) UMAP embedding from a human MASLD liver snRNA-seq dataset, colored by annotated cell type (A) or disease stage (B). (C) Dotplot showing expression of the 39 prioritized candidate genes across liver cell types; columns correspond to cell types and rows to genes. Dot size indicates the percentage of cells expressing each gene, and color encodes scaled average expression (red, higher; blue, lower).

**Figure S6.**
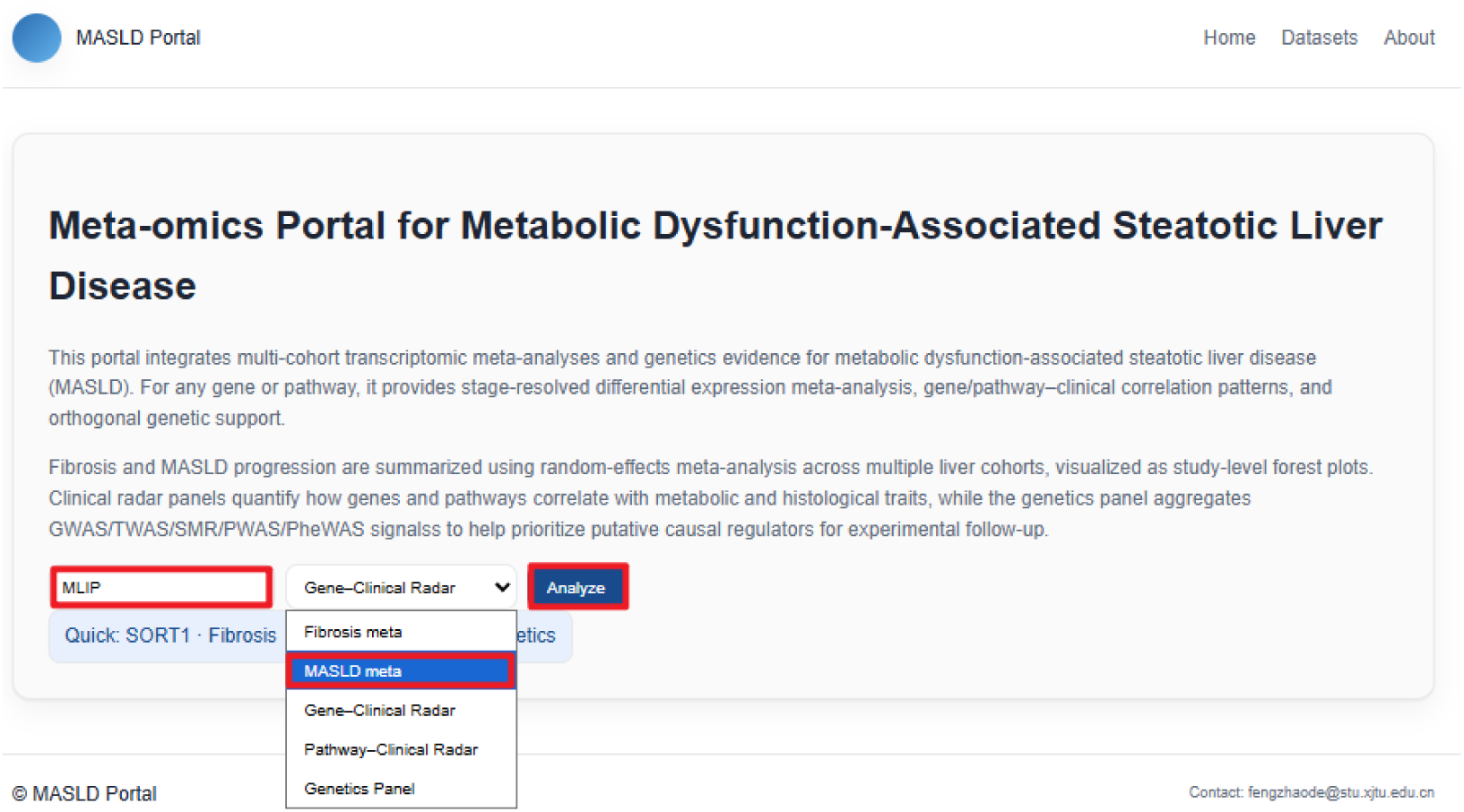
Overview of the MASLD Portal. Screenshot of the web interface enabling gene-centric queries and selection of analytic views, including MASLD stage meta-analysis, fibrosis-stage meta-analysis, gene–clinical correlation, pathway–clinical correlation, and an integrated genetics panel. The portal provides interactive visualization of multi-cohort transcriptomic meta-analysis and orthogonal genetic evidence to facilitate exploration and prioritization of candidate MASLD regulators.

