## Supplemental Table 16 for "Integrative multi-omics and multi-trait analysis prioritizes regulatory mechanisms and genes for metabolic dysfunction-associated steatotic liver disease"

|  | **Primer sequence** | |
| --- | --- | --- |
| **Gene** | **Forward** | **Reverse** |
| **β-actin** | CATGTACGTTGCTATCCAGGC | CTCCTTAATGTCACGCACGAT |
| ***MLIP*** | TTATTGTGGACTCCGAAGGGG | GCACTACATCAGAGACCAGACTG |

Supplementary Table 16. Sequences of primers used for real-time quantitative PCR.
